# The environment selects: Modeling energy allocation in microbial communities under dynamic environments

**DOI:** 10.1101/689174

**Authors:** Leonor Guedes da Silva, Sergio Tomás-Martínez, Mark C.M. van Loosdrecht, S. Aljoscha Wahl

## Abstract

What will be the best metabolic strategy in a competitive environment where oxygen is periodically unavailable? A few decades ago, an accidental, man-made cyclic anaerobic/aerobic environment selected for Polyphosphate Accumulating Organisms (PAOs) and this strategy is now widely used to allow for Enhanced Biological Phosphorus Removal (EBPR) of wastewater. But could it have been predicted? Here, a dynamic resource allocation modeling formalism was used to analyze the impact of selection pressures on metabolic function. With the same meta-network but modified selective pressures, different successful strategies can be predicted: Polyphosphate-AOs, Glycogen-AOs, Polyhydroxyalkanoate-AOs, and regular aerobic heterotrophs. The results demonstrate how storage metabolism allows for different trade-offs between growth yield, robustness, and competitiveness, and highlight how each metabolic function is an important determining factor for a selective advantage in a given environment. This can be seen as an example of when *“Unity in biochemistry”* by A.Kluyver meets *“Everything is everywhere, but the environment selects”* by B.Becking and how microbial ecosystems may be described by the energy allocation phenotype instead of a detailed description of each organism.

## INTRODUCTION

About 85 years ago, Baas Becking hypothesized that all microbial life is distributed worldwide, but the environment (e.g. gut, skin, plants, wastewater treatment systems) selects for and enhances a specific phenotypic function, which then becomes observable (i.e. above detection limit) (Baas-Becking, 1934). To date, different functions in microbial ecosystems have been described using computational models (reviewed in (Succurro & Ebenhöh, 2018)), however predictive power is still limited. For example, current kinetic models use pre-defined optimal metabolic strategies and require extensive re-calibration to simulate different environments. Consequently, these models cannot predict the outcome of a new environmental condition.

Natural environments are dynamic and changing conditions are inevitable. To cope with this, evolution has selected for a certain degree of metabolic flexibility that leads to the recently discussed rate/yield trade-offs (Abudukelimu, Mondeel, Barberis, & Westerhoff, 2017; Frank, 2010; Peyraud, Cottret, Marmiesse, Gouzy, & Genin, 2016; Pfeiffer, Schuster, & Bonhoeffer, 2001). Depending on the environmental pressure, different cellular characteristics arise as selective advantage, e.g. higher substrate uptake or growth rates *versus* higher energetic efficiency (i.e. higher biomass yield) *versus* membrane space (i.e. higher transporter capacity) *versus* storage metabolism (i.e. metabolic buffer capacity).

Proposed solutions to tackle some of today’s grand societal challenges like wastewater treatment as well as production of chemicals from renewable resources or waste streams rely on microbial communities. In the design of these processes, engineers can exploit the selective advantage of a given community (i.e. intracellular storage) to, for example, remove phosphate or produce polyhydroxyalkanoates (PHA) from wastewater streams (Barnard, 1976; Kleerebezem & van Loosdrecht, 2007). To select for a relevant microbial community, one of the main technological questions arising is “how to design and control the selective environment that stabilizes the open microbial community for the desired function?”.

One of the best characterized and modeled microbial communities can be found in biological wastewater treatment plants. The selected microbial community was tuned over the years to promote removal of organic carbon, nitrogen and phosphorus from sewage. In its history, there have been several discoveries which led to the design of new generations of this bioprocess. An important condition for enhanced biological phosphorus removal (EBPR) was accidentally discovered in treatment plants that contained an anaerobic zone at the entrance in otherwise fully aerobic activated sludge systems (Srinath, Sastry, & Pillai, 1959). As a result, microorganisms experienced time-varying presence and absence of external electron acceptors (O_2_). In the absence of O_2_, ordinary aerobic heterotrophs cannot produce energy, which gives a selective advantage to Polyphosphate Accumulating Organisms (PAOs) that can use their polyphosphate and glycogen storage for energy production. PAOs remove phosphate from the environment by accumulating it intracellularly as polyphosphate (Barnard, 1976; Seviour, Mino, & Onuki, 2003).

When no external electron acceptor is present (usually defined as anaerobic condition in EBPR literature), PAOs use their polyphosphate and glycogen storage as competitive advantage (Van Loosdrecht, Pot, & Heijnen, 1997). These storage compounds are used to generate ATP and NADH allowing to rapidly sequester extracellular organic carbon sources such as volatile fatty acids to store them intracellularly as polyhydroxyalkanoates (PHAs). The incorporation of these sources into PHAs is faster than into biomass synthesis (growth), representing another competitive advantage: rapid sequestration of the extracellular organic carbon sources making these inaccessible for the regular heterotrophs. In the presence of an external electron acceptor (aerobic), the accumulated PHAs are used for both growth and regeneration of the polyphosphate and glycogen storage pools. While the single metabolic traits of PAO are commonly found among microorganisms (genotype), the combined use of them to sequester substrates in the absence of external electron acceptors is what defines the PAO phenotype. For example, in periodic, dynamic environments such as activated sludge systems, other microbial functional groups can be found such as Glycogen Accumulating Organisms (GAOs), Polyhydroxyalkanoate Accumulating Organisms (PHA-AOs) and regular aerobic heterotrophs (Figure 1); these microorganisms share the single metabolic traits of PAOs, however each uses them to different extents (phenotype) which result in different levels of fitness (i.e. reproductive success in a given environment). An additional investment of resources in these storage phenotypes confers microorganisms robustness to cope with unexpected events during the cyclic environment (Maurer, Gujer, Hany, & Bachmann, 1997). Such additional storage activity requires resources (i.e. metabolic energy, but also additional enzymes/proteins) leading to a reduction of the maximal growth rate and yield compared to a growth-only strategy and generates a trade-off between the different metabolic processes (Figure 1).

**Figure 1.**
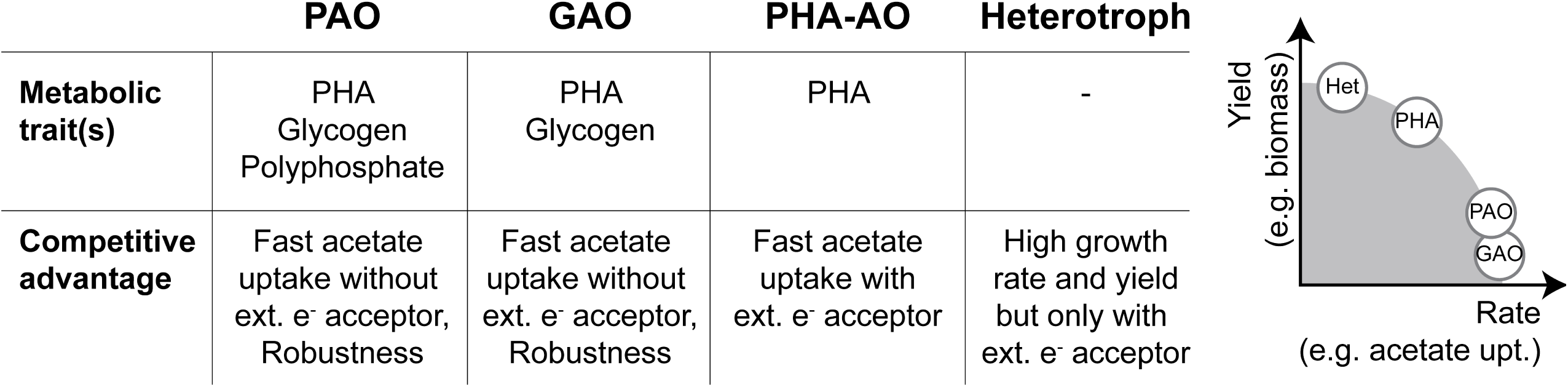
Main phenotypes found in activated sludge systems. Left: Metabolic traits and their respective competitive advantages. Right: Expected rate/yield trade-off curve.

Recently, a modeling approach for dynamic resource allocation has been proposed that integrates stoichiometry and dynamic conditions and allows for the calculation of optimal phenotypes. This new modeling framework (Rügen, Bockmayr, & Steuer, 2015) known as Conditional Flux Balance Analysis (cFBA) is a variant of dynamic FBA and it has been developed as a dynamic resource allocation formalism to understand growth in a periodic environment. This formalism has been used previously for the analysis of axenic cyanobacterial growth in day/night cycles (Faizi, Zavřel, Loureiro, Červený, & Steuer, 2018; Reimers, Knoop, Bockmayr, & Steuer, 2017; Rügen et al., 2015).

In the present study we aim to better understand why each different metabolic strategy – PAOs, GAOs, PHA-AOs and heterotrophs – despite their common genetic traits, can all be found in a competitive environment where oxygen is periodically unavailable. To do so, we constructed a model using the novel cFBA approach (Figure 2) to:

**Figure 2.**
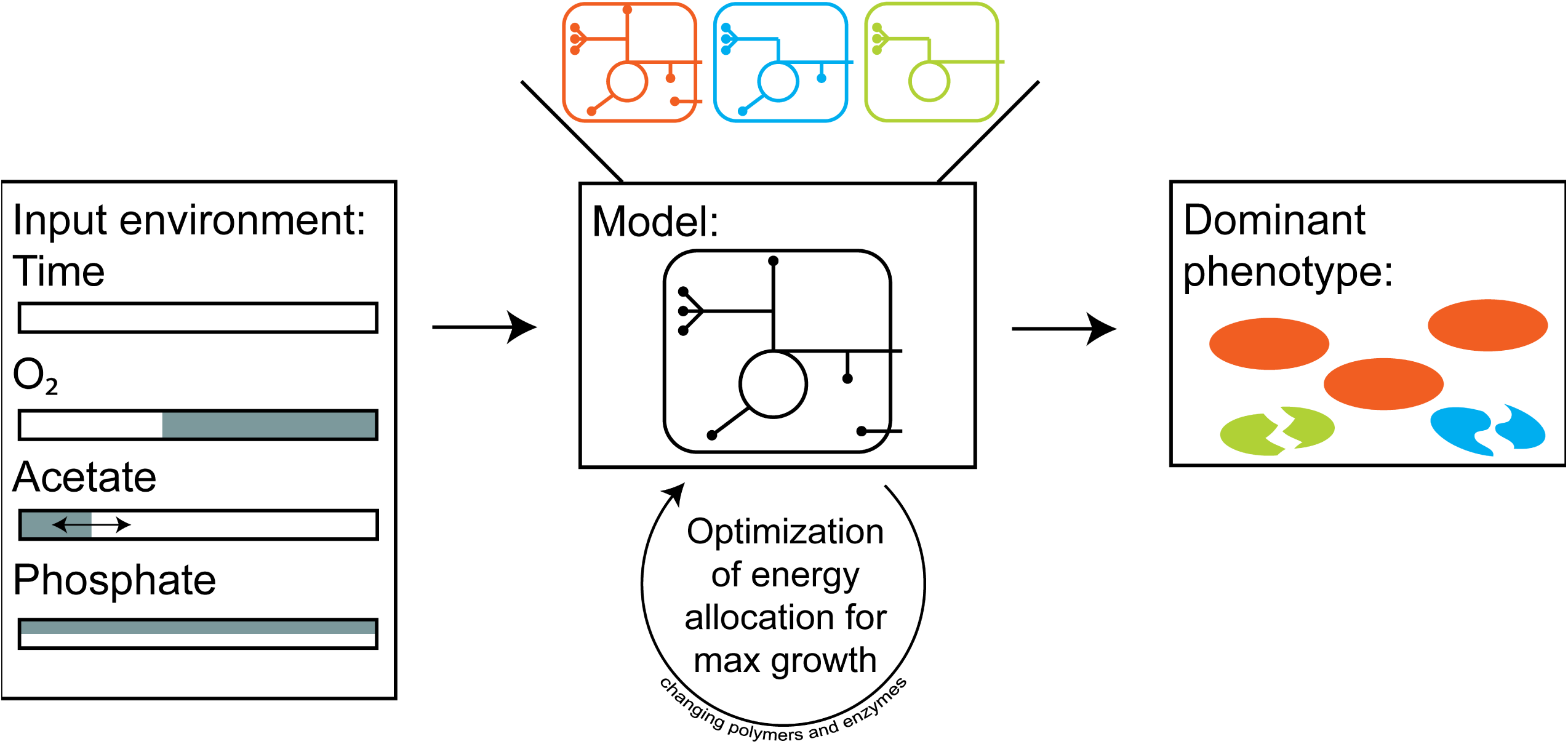
Overview of the approach used in this study. Cycle time and anaerobic/aerobic phases are fixed, while acetate uptake period and external phosphate availability are varied to create different environments.

1. Demonstrate that investment in (polyphosphate and/or glycogen) storage leads to improved fitness, generating a competitive advantage as well as robustness under periodic anaerobic/aerobic conditions;
2. Compare the investments on the two alternative polyphosphate or glycogen-based metabolic (PAM or GAM) strategies and show how PAOs can still adapt their metabolic strategy when polyphosphate storage is limiting;
3. Explore how the environment selects for the phenotype (metabolic trait) with highest fitness and how that leads to different rate/yield trade-offs.

## MODEL CONSTRUCTION

In the framework of cFBA the cell is described as an autocatalytic system with different functions that contribute to growth and share common resources. Maximal reaction rates are defined by the amount of available enzyme and its respective catalytic capacity. Consequently, enzyme synthesis is also limited by the quantity of ribosomes, which in turn are synthesized by enzymes (auto-catalysis).

Here, the amounts of macromolecules such as biomass precursors, proteins, enzymes, ribosomes and storage polymers are explicitly modeled and are considered time-dependent (i.e. dynamic). Moreover, alike in conventional FBA, it is assumed that changes in metabolic intermediates occur much faster than in macromolecules, thus all intermediates are in *quasi-steady* state. To enforce the synthesis of inert compounds like biomass precursors and non-catalytic proteins, a so-called quota (i.e. minimum amount required) is introduced (Rügen et al., 2015). Quotas for the storage polymers are also set to represent the extra investment in robustness.

In this framework, dynamic environmental transitions are described as stable, periodic cycles. Consequentially, biomass composition is the same at the end and start of a cycle. Furthermore, when such repetitive system is stable, the biomass synthesized during one cycle equals the biomass removed from the system (i.e. in equals out). Based on that, the net growth at end of the cycle is expressed as a multiplication of the defined individual cellular components at the beginning of the cycle.

All simulations were based on a simplified metabolic network of one of the most well studied PAOs, *Candidatus* Accumulibacter phosphatis, derived from (Oyserman, Noguera, del Rio, Tringe, & McMahon, 2016). This model was constructed following the steps proposed by (Reimers, Lindhorst, & Waldherr, 2017).

### Metabolic network

The model comprises the main metabolic reactions to describe PAOs growth during anaerobic-aerobic cycles (Figure 3). This works as a meta-network since the phenotypes of GAOs, PHA-AOs and heterotrophs are subsets of the metabolic network of PAOs. Traditionally metabolic models for these organisms have been separated for the different redox conditions, the here used model apprch does not need predefined modelling of anaerobic or aerobic conditons. Reaction names and stoichiometry are shown in Table 1. ATP requirements for acetate and phosphate uptake, together with the ATP:NADH stoichiometry of the oxidative respiration (i.e. P/O ratio in ETC reaction) were based on (Smolders, van der Meij, van Loosdrecht, & Heijnen, 1994a, 1994b, 1995). Phosphoenolpyruvate (PEP) is introduced as a link between catabolism and biosynthesis. As simplification, the different biomass precursors (BMP) are modeled as one quota component. This metabolite represents components of biomass such as DNA, RNA and phospholipids. The stoichiometry for the anabolic reactions was based on (Oehmen, Zeng, Keller, & Yuan, 2007) for BMP, and (Rügen et al., 2015) for ribosomes and proteins. The different types of PHA polymers (PHB, PHV, PH2MV, PH2MB) originating from different combinations of the precursors Acetyl-CoA and Succinyl-CoA were simplified as PHB and PH2MV. With this simplification, 1 monomer of PHV or PH2MB is described by half a molecule of PHB and half a molecule of PH2MV.

**Table 1.**
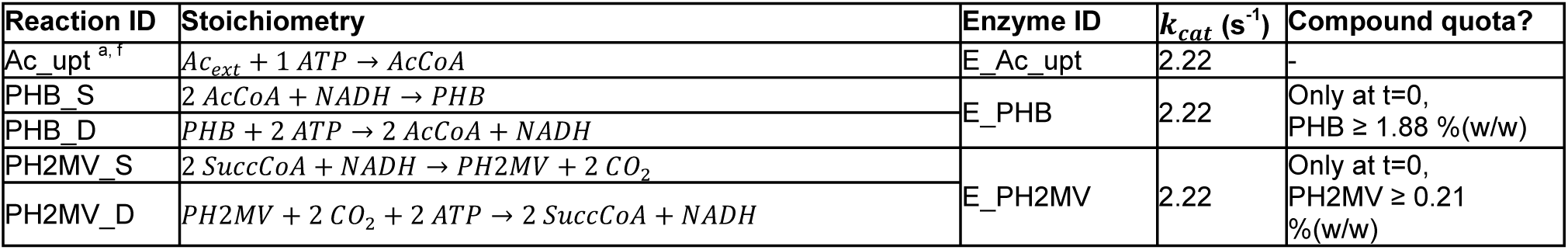
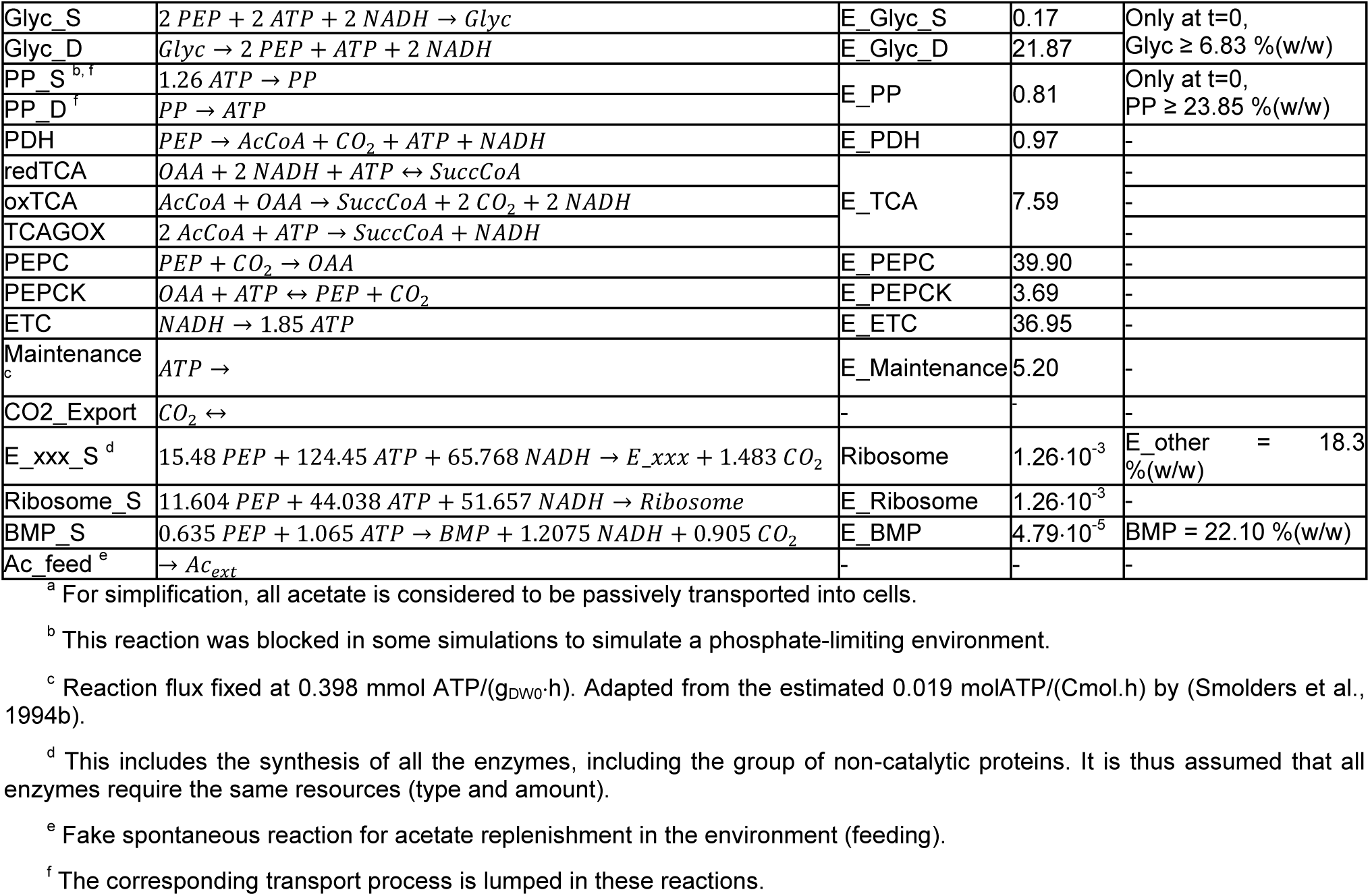
Complete reaction network, quotas and turnover rates used in this model.

**Figure 3.**
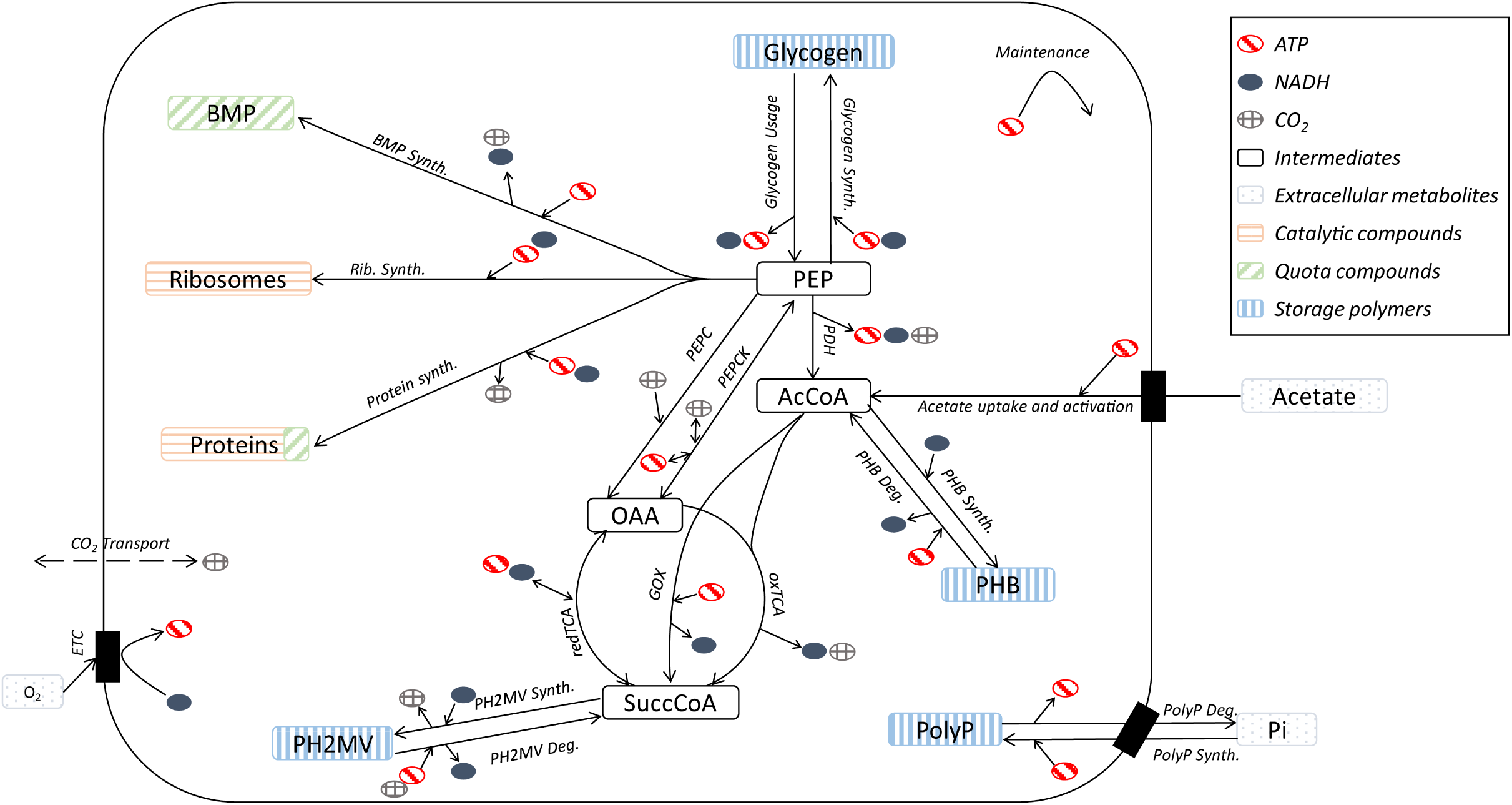
Simplified representation of the metabolic network used in this study. The complete reaction list and respective stoichiometries are shown in Table 1.

A maintenance reaction is introduced to represent energy requirements of the cellular processes like protein turnover among others, which currently can only be estimated from experimental data, here taken from (Smolders et al., 1994b). Lastly, all redox cofactors are expressed as NADH, and all energy cofactors as ATP.

### Biomass composition, quotas and enzyme capacities

In this model, biomass is composed of storage polymers and lean biomass (sum of all proteins, ribosomes, and BMP). In order to ensure the synthesis of BMP and non-catalytic proteins, minimum amounts (quotas) for those compounds are defined for all time points. On the other hand, for storage polymers (glycogen, polyphosphate and PHA), only an initial quota is needed. These values were derived from literature data: a) for glycogen, polyphosphate and PHAs, experimental measurements from (Acevedo et al., 2012) were used, b) total protein content was based on the work of (Yücesoy, Lüdemann, Lucas, Tan, & Denecke, 2012), c) composition of this protein pool was based on the metaproteomic study by (Barr et al., 2016), and d) for the ribosome content, no PAO data was available – here the value used in (Rügen et al., 2015) for cyanobacteria was taken. An overview of quota values for these compounds can be found in Table 1.

Except for CO_2_ diffusion, all reactions in the system are catalyzed by enzymes or ribosomes. Constraints for the (maximal) biomass specific reaction rates are defined by the catalyst amounts and their specific activities (represented by the turnover rates, *k*_*cat*_). Rügen and co-workers estimated *k*_*cat*_ values for each modeled reaction (often representative of whole pathways) and assuming all enzymes have the same production cost (Rügen et al., 2015). For reactions missing in their model, the turnover rates were derived from the work of Davidi and colleagues (Davidi et al., 2016) by scaling them based on common reactions presented in both studies. Since these values were estimated for cyanobacteria and *E. coli*, respectively, it is reasonable to also use them to describe the bacterial ecosystems understudy (see Table 1). Furthermore, a sensitivity analysis was performed to study the influence of these values, i.e. each *k*_*cat*_ parameter was changed by one order of magnitude up and down (see Figure 8 in Supplementary Information).

**Figure 4.**
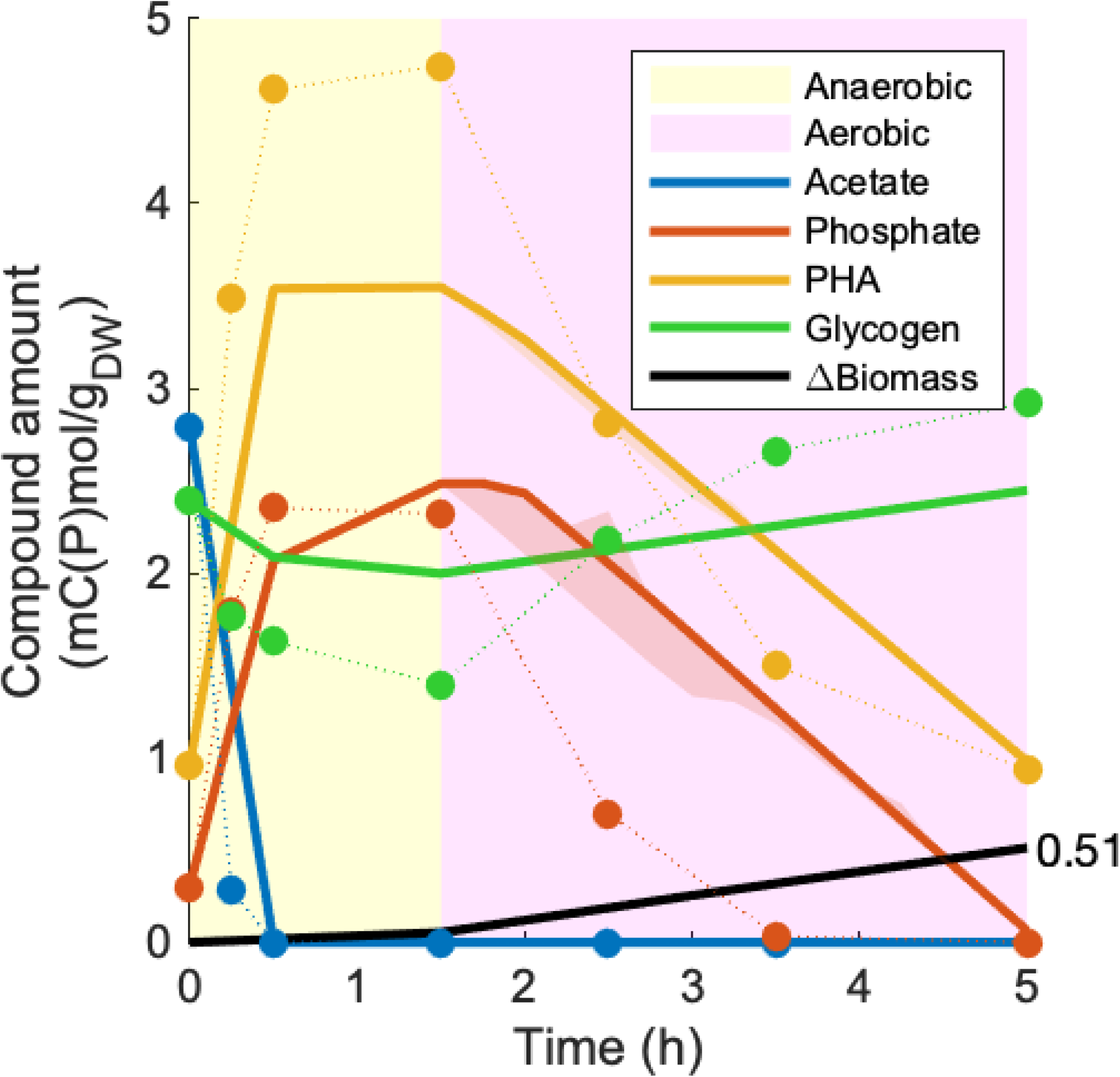
Simulated and experimental amounts of extracellular acetate and phosphate, intracellular polymers (PHA, glycogen) and lean biomass. Amounts are normalized to the initial biomass amount. The phosphate profile is calculated from the polyphosphate dynamics and initial phosphate supplied in the medium (experimental). The simulated PHA line is calculated from the modeled PHB and PH2MV pools. Solid lines represent the simulation and bullets represent experimental data retrieved from Fig. 3 of (Acevedo et al., 2012).

**Figure 5.**
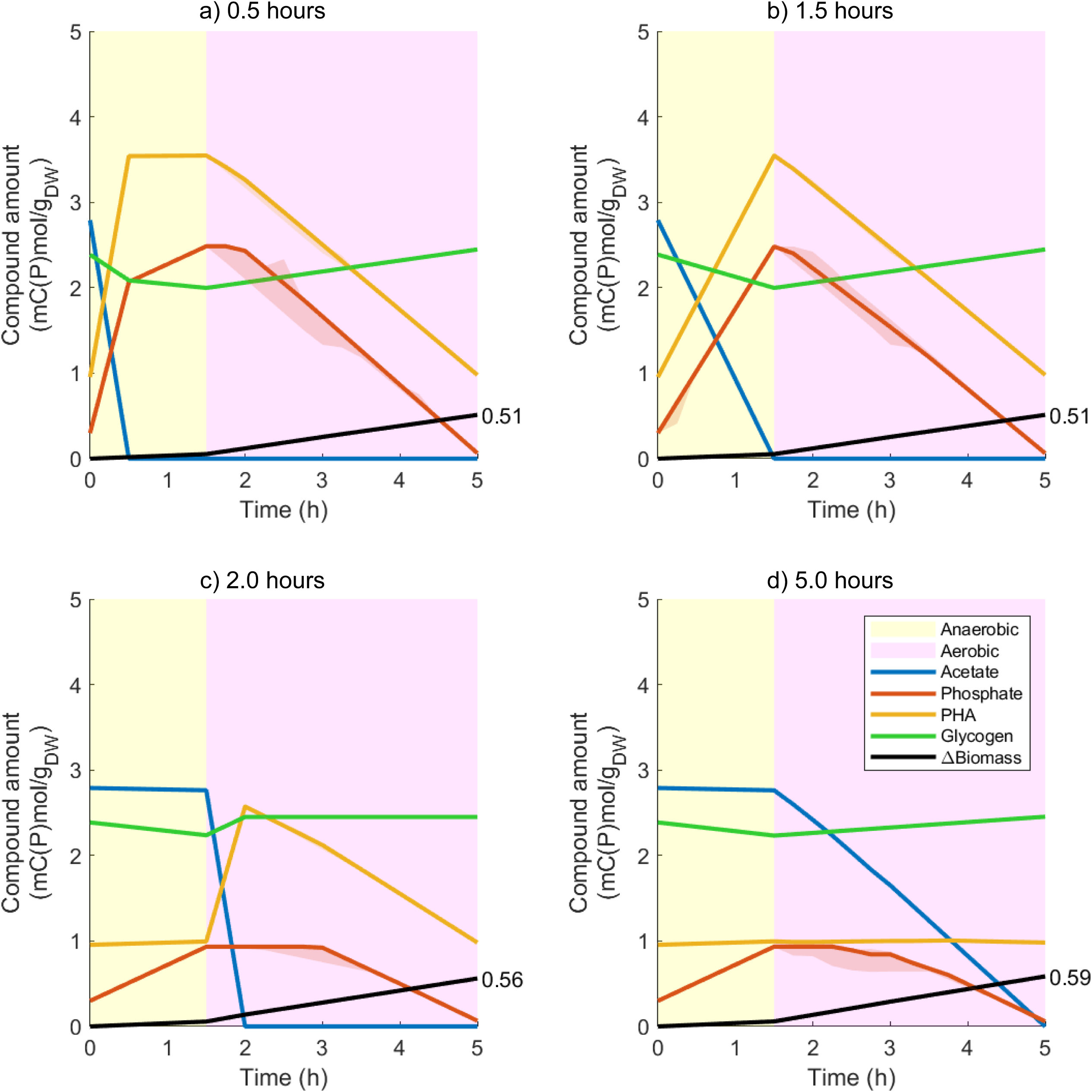
Simulated amounts of extracellular acetate and phosphate, intracellular polymers (PHA, glycogen) and lean biomass. The shaded areas around the simulated lines represent the output of the flux variability analysis. Amounts are normalized to the initial biomass amount. The normalized amount of new biomass at the end of the cycle is explicitly shown. The phosphate profile is calculated from the polyphosphate dynamics and initial phosphate supplied in the medium (experimental). The simulated PHA line is calculated from the modeled PHB and PH2MV pools. Scenarios: a) acetate uptake set to last only 30 min (same as in Figure 4); b) 1.5h; c) 2h and d) 5h.

**Figure 6.**
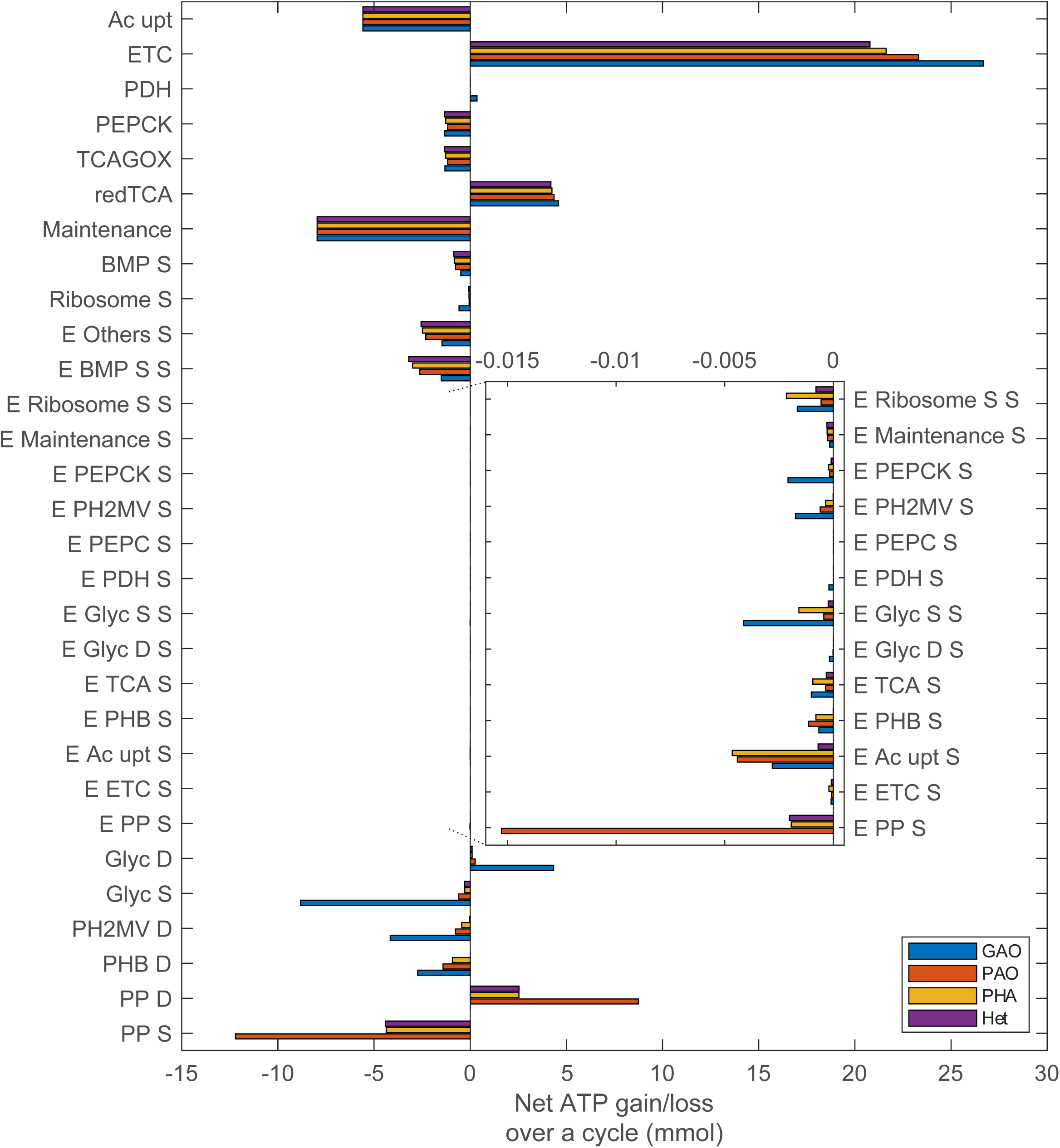
Net ATP gain/loss over a cycle for each different cellular reaction for each metabolic strategy (excluding the ATP cost of producing the enzyme catalyzing the reaction). For each storage polymer the synthesis (S) reaction is separated from the degradation (D) reaction. The zoomed inset shows the ATP costs of synthesizing each enzyme (E).

**Figure 7.**
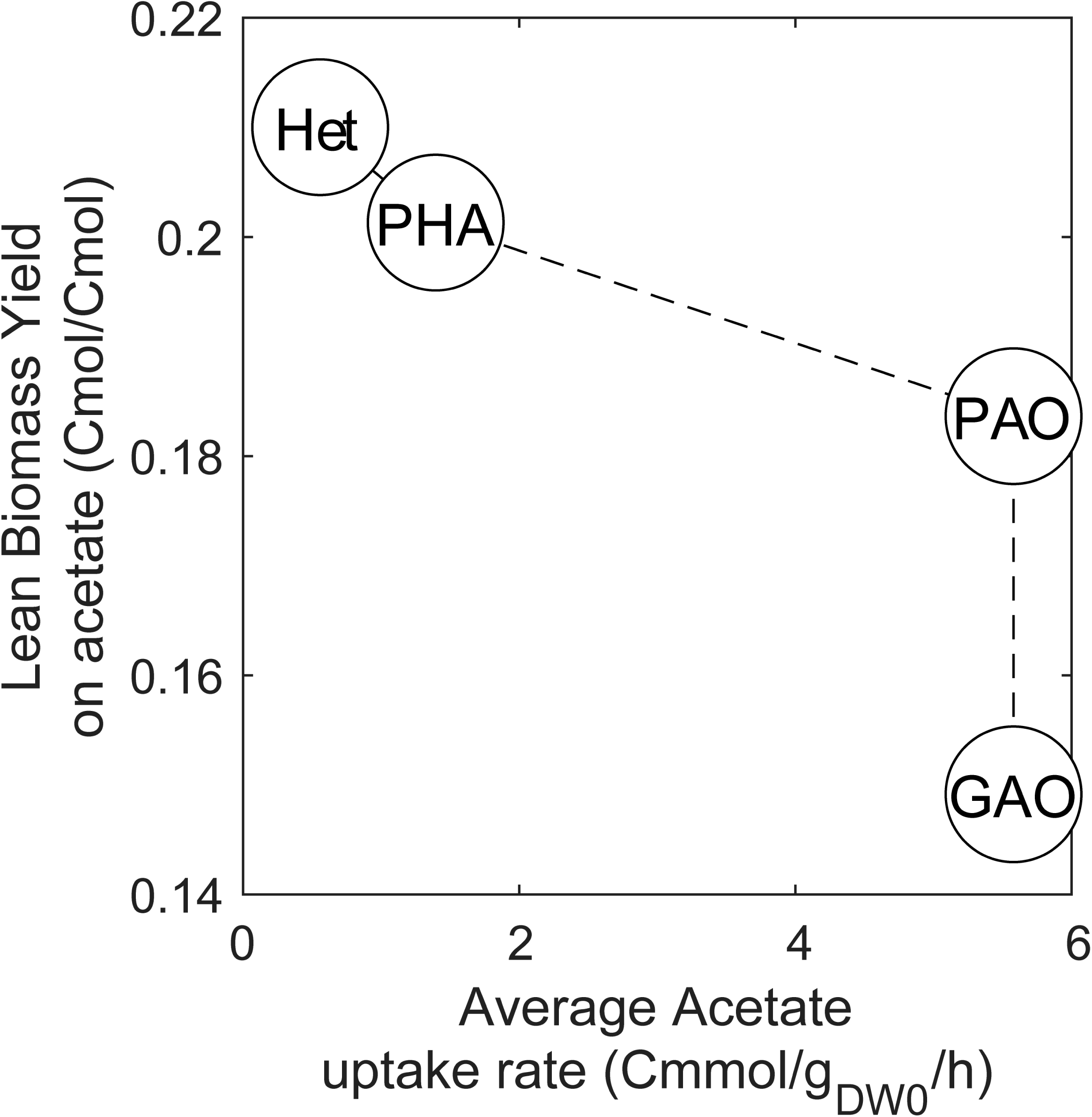
Rate/Yield trade-off curve. The curve was obtained based on the simulated results of the different metabolic strategies shown in Figure 5 and Table 2.

**Figure 8.**
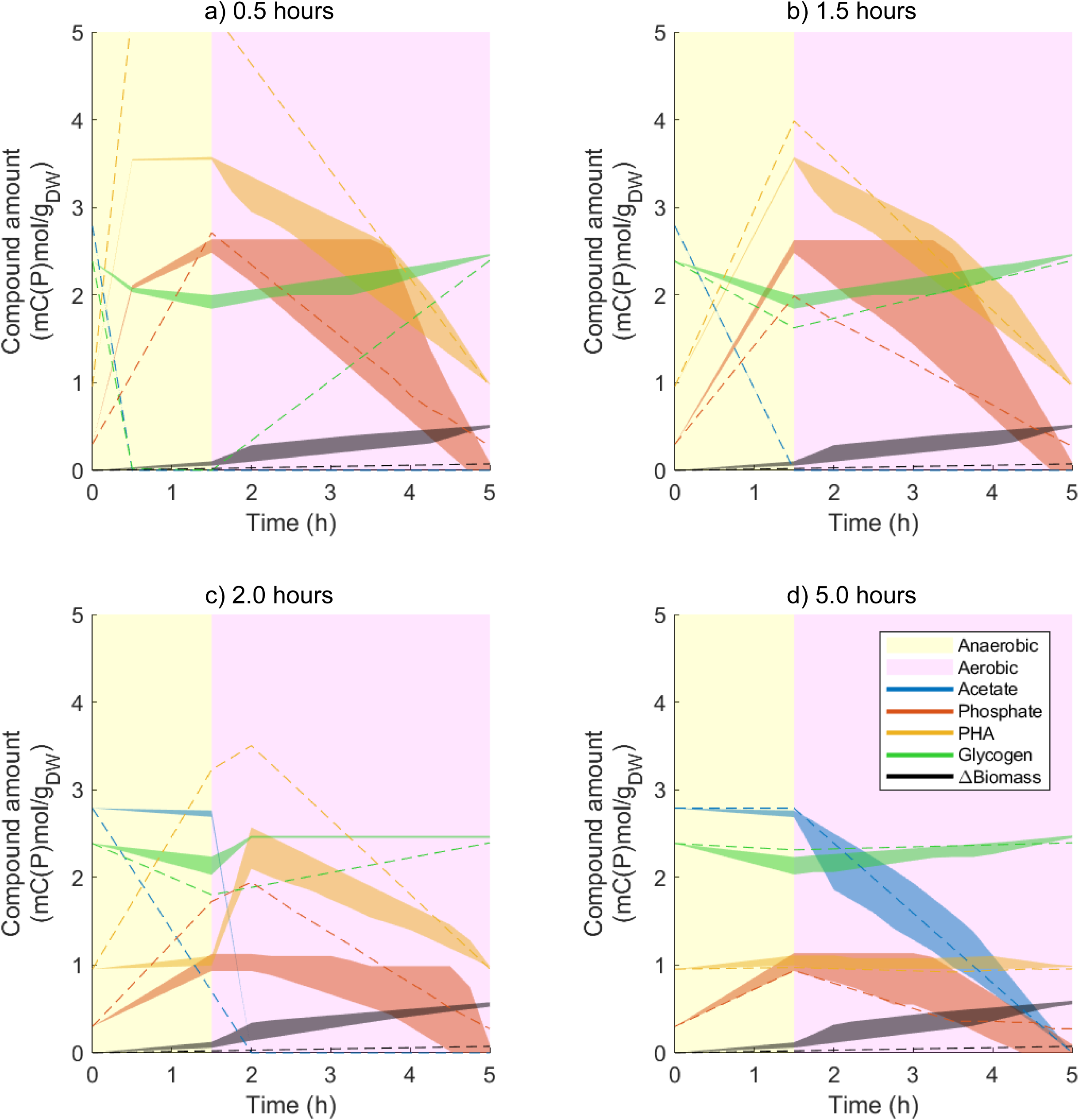
Simulated amounts of extracellular acetate and phosphate, intracellular polymers (PHA, glycogen) and lean biomass for the sensitivity analysis of the turnover rates (*k*_*cat*_). Individual values of the different *k*_*cat*_ were changed by a factor of 0.1 and 10. Shaded areas represent the grouped output of all the simulations, except for the decrease of *k*_*cat*_ for BMP synthesis, which is represented by the dashed line. Amounts are normalized to the initial biomass amount. The phosphate profile is calculated from the polyphosphate dynamics and initial phosphate supplied in the medium (experimental). PHA is calculated from PHB and PH2MV. Scenarios: a) acetate uptake set to last only 30 min (same as in Figure 4); b) 1.5h; c) 2h and d) 5h.

## RESULTS AND DISCUSSION

A competitive environment where oxygen is periodically unavailable was simulated using the cFBA model as previously described. As reference simulation, the experimental conditions (i.e. initial amounts of extracellular acetate, phosphate and intracellular polymers, and the duration of the phases) presented by (Acevedo et al., 2012) were chosen. The authors reported a high enrichment of PAOs in their reactors based on high phosphate release upon acetate uptake and visual confirmation by FISH.

The reference simulation reproduced known features of PAOs’ metabolism (Figure 4) such as:

- Polyphosphate utilization as energy source for anaerobic acetate uptake;
- Glycogen degradation provides reducing equivalents anaerobically and extra ATP;
- Anaerobic PHA accumulation;
- Aerobic PHA degradation for biomass, glycogen and polyphosphate synthesis;

Apart from the minimal quotas and turnover rates as mentioned earlier, only 3 additional constraints were needed to simulate this PAO phenotype: (1) amount of intracellular storage polymers at the beginning of the cycle, (2) amount of acetate available extracellularly per cycle and (3) maximum amount of time cells have to completely consume it (feast duration). The combination of (2) and (3) sets the minimum acetate uptake rate. The type of metabolic response was not a constraint in this model, nor any of the other kinetic rates.

While the amount of acetate per g_DW0_ available is an experimental design input defining the environment, the constraints on storage quotas are specific to PAO metabolism. When no initial amount of intracellular storage polymers is set, the simulation predicts that the (optimal) cell will only accumulate the minimum amount of polymers needed to consume all acetate anaerobically; polyphosphate and glycogen would optimally be zero at the anaerobic/aerobic switch, and PHA would be zero at the beginning/end of the cycle. These are specific optima for the defined environmental conditions, and, as consequence, these cells have no buffer capacity to cope with fluctuations in the environment. Investing in higher accumulation as observed experimentally will increase the robustness during a deviating cycle as reserves will be available. The initial intracellular storage polymer quotas thus represent a growth trade-off towards robustness.

Furthermore, the acetate uptake duration was constrained to obtain the PAO phenotype. In a simulation without such constraint, acetate is consumed only during the aerobic phase (as seen later in Figure 5). This constraint thus selects for specific phenotypes with respect to a rate/yield tradeoff. Substrate uptake rate is an essential competitive advantage, as the fastest consuming organism will thrive. Anaerobic acetate consumption represents a significant rate advantage (i.e. earlier sequestration) compared to waiting for aerobic conditions. However, and as later seen in Figure 6, this represents a trade-off against growth as a significant allocation of energy is required for formation and consumption of the stored PHA. In this case, the allocated energy is mainly used for polymer cycling instead of the traditional protein resource allocation used to describe e.g. Crabtree effect in yeast (Nilsson & Nielsen, 2016).

### Polyphosphate *versus* glycogen

The reference simulation representing the optimal operation for the given condition (Figure 4) deviates from the experimental observations for the glycogen usage (Acevedo et al., 2012). In their experiment as well as in the work of Welles and colleagues (Welles et al., 2015), a metabolic shift of PAOs from a polyphosphate-based metabolism (PAM) to a glycogen-based metabolism (GAM) was associated with different experimental conditions. This metabolic flexibility was also discussed extensively from a redox balancing viewpoint where different sources of reducing equivalents were considered: partial TCA cycle with/without glyoxylate shunt, and glycolysis (Silva et al., 2018). One described environmental condition that potentially promotes GAM is low phosphate availability. To simulate this limitation, the current model was used, but this time with polyphosphate synthesis impeded. As a result of this simulation, a GAM phenotype came up as the optimal strategy (Table 2). Note that GAM is also the strategy used by GAOs.

**Table 2.**
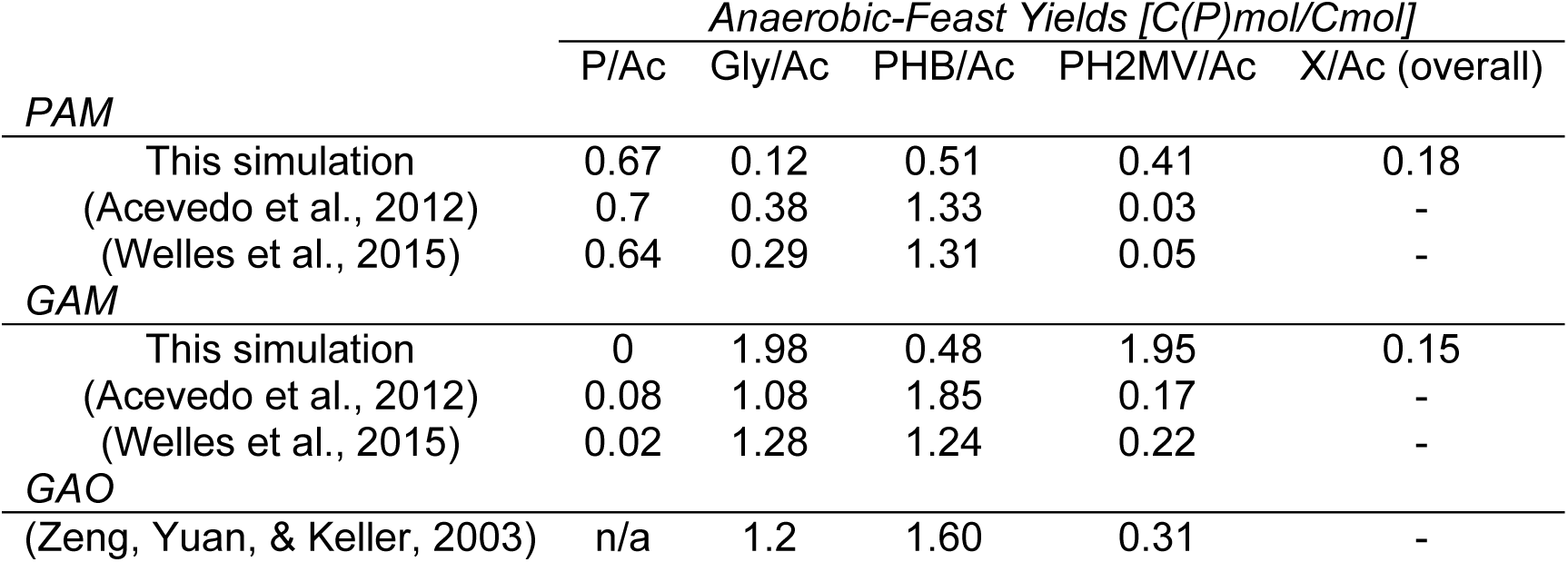
Simulated and experimental anaerobic-feast yields [C(P)mol/Cmol]. Feast corresponds to the period in which acetate is present. Results are shown for PAM (PAOs performing polyphosphate-based metabolism), GAM (PAOs performing glycogen-based metabolism) and GAOs. Experimental values were converted so that 1 monomer of PHV (or PH2MB) is described by half a molecule of PHB and half a molecule of PH2MV.

|  | *Anaerobic-Feast Yields [C(P)mol/Cmol]* |  |  |  |  |
| --- | --- | --- | --- | --- | --- |
|  | P/Ac | Gly/Ac | PHB/Ac | PH2MV/Ac | X/Ac (overall) |
| *PAM* |  |  |  |  |  |
| This simulation | 0.67 | 0.12 | 0.51 | 0.41 | 0.18 |
| (Acevedo et al., 2012) | 0.7 | 0.38 | 1.33 | 0.03 | - |
| (Welles et al., 2015) | 0.64 | 0.29 | 1.31 | 0.05 | - |
| *GAM* |  |  |  |  |  |
| This simulation | 0 | 1.98 | 0.48 | 1.95 | 0.15 |
| (Acevedo et al., 2012) | 0.08 | 1.08 | 1.85 | 0.17 | - |
| (Welles et al., 2015) | 0.02 | 1.28 | 1.24 | 0.22 | - |
| *GAO* |  |  |  |  |  |
| (Zeng, Yuan, & Keller, 2003) | n/a | 1.2 | 1.60 | 0.31 | - |

The purpose of utilizing glycogen or polyphosphate anaerobically is the same: to provide ATP for anaerobic acetate uptake and activation for polymerization into PHAs. The choice of one ATP source over the other can be determined by the environment. As shown in Table 2, PAM leads to a higher growth yield than GAM. This is because polyphosphate production and consumption are energetically less costly than the glycogen cycling (Figure 6). Furthermore, glycogen cycling comes along with balancing reducing equivalents whereas polyphosphate not. As demonstrated in (Silva et al., 2018), polyphosphate leads to higher metabolic flexibility thus allowing for higher fitness to cope with changes in the environment. However, when phosphate is limited in the environment and polyphosphate accumulation is impeded, GAM is the alternative. In both PAM and GAM simulations, the acetate uptake rate and the ATP costs for both acetate and phosphate transport were set to the same value. However, these three parameters change differently for PAOs and GAOs under different environmental conditions such as pH and temperature (Lopez-Vazquez et al., 2009). Consequentially, each parameter combination may select for either PAM or GAM, as a lower biomass yield can be compensated with a higher acetate uptake rate.

### “…but the environment selects”

There are critical environmental characteristics that dictate the most favorable metabolic strategy (trait), e.g. rate-*versus* yield-strategy (Maharjan et al., 2013). Here, different rate/yield trade-offs are further analyzed by constraining the maximal time for acetate uptake during the cyclic feeding regime (rate-selective pressure). Based on the predicted optimal metabolic function, the dominance of PAOs/GAOs/PHA-AOs/heterotroph’s strategy is discussed.

While keeping the same core model, the maximal time for acetate uptake was varied from 30 minutes (reference condition described by (Acevedo et al., 2012)) up to 5 hours (the whole cycle time). The simulated scenarios show distinct metabolic strategies (Figure 5).

When acetate is allowed to be consumed throughout the whole cycle (Figure 5d), the optimal solution found is to only do it once oxygen is available and only regular aerobic heterotrophic growth occurs. On the other hand, when acetate is forced to be taken up within 30 min into the anaerobic phase (Figure 5a), the optimal solution is to invest in the storage of polyphosphate and glycogen aerobically and to make use of them anaerobically. As it can be seen from the amount of lean biomass synthesized in each case, storing polyphosphate and glycogen is indeed an investment as it leads to a lower biomass yield (Figure 5a, 0.18 Cmol biomass per Cmol acetate) as opposed to the case where no storage metabolism is needed (Figure 5d, 0.21 Cmol biomass per Cmol acetate). These biomass yields on acetate obtained in the simulations are comparable with the 0.24 Cmol biomass per Cmol acetate reported by (Acevedo et al., 2012) for the experimental conditions here simulated.

Interestingly, the amount of biomass produced is not as sensitive to the anaerobic acetate uptake rate (0.5 and 1.5h scenarios), as opposed to when acetate is consumed fully aerobically (2 and 5h). If the anaerobic biomass yields are the same, then the fastest consumer will thrive. This raises the question: what is then the bottleneck setting the maximal biological rates in these systems? Here it is already seen that PHA production alleviates the growth bottleneck allowing for faster acetate uptake. Thus, the question remains whether PHA production has enough processing capacity to keep up with acetate transport, or if this transport is the actual bottleneck. Other possible bottlenecks may be related with the ATP generation capacity or any other related process such as limited membrane space for transporters. To shed light on these questions, we recommend a comprehensive analysis of biomass composition, proteome (incl. transporters), membrane structure and processes.

### Intracellular energy allocation strategies and respective rate/yield trade-offs

To further explore the investment on each cellular process, their net ATP gain/losses are shown in Figure 6. At first glance, the differences in ATP generation in the ETC reaction point towards the efficiency of the strategy employed: Less ATP invested means less need for its production, and thus leading to a higher biomass yield as verified earlier.

Commonly, in rate/yield trade-off analyses, the costs of higher rates (enzymes) are the culprit for the biomass yield loss caused by such investment. For the accumulating organisms (AOs) under study, there is indeed an extra investment on enzymes to enable the observed polymer cycling. However, this energy requirement is about 100 times lower (see zoomed section of Figure 6) than the ATP involved in the polymer cycling itself (i.e. ATP requirements for transport and activation reactions). One could also argue on the investment in different enzymes, however, this model is too coarse in that respect. For a more detailed analysis on protein allocation, this model requires further improvements as it currently assumes a) all enzymes have the same production cost and b) global catalytic capacities *(k*_*cat*_) are estimated for each linear pathway, which are currently based on *in vitro* measurements but should become predictive using thermodynamics and molecular dynamics modeling (Pekař, 2015). Nonetheless, a sensitivity analysis was made by varying each assumed *k*_*cat*_ one order of magnitude higher/lower (see Figure 8 in Supplementary Information). The results show that the simulated output is only very sensitive to the 10x decrease of *k*_*cat*_ for BMP synthesis. However, the simulated biomass yield in that case is one order of magnitude lower than the experimentally observed, which is further from reality than the one obtained in our reference simulation. For all the other simulations, all conclusions were not sensitive to changes in *(k*_*cat*_) across 3 orders of magnitude (see Figure 9 in Supplementary Information).

**Figure 9.**
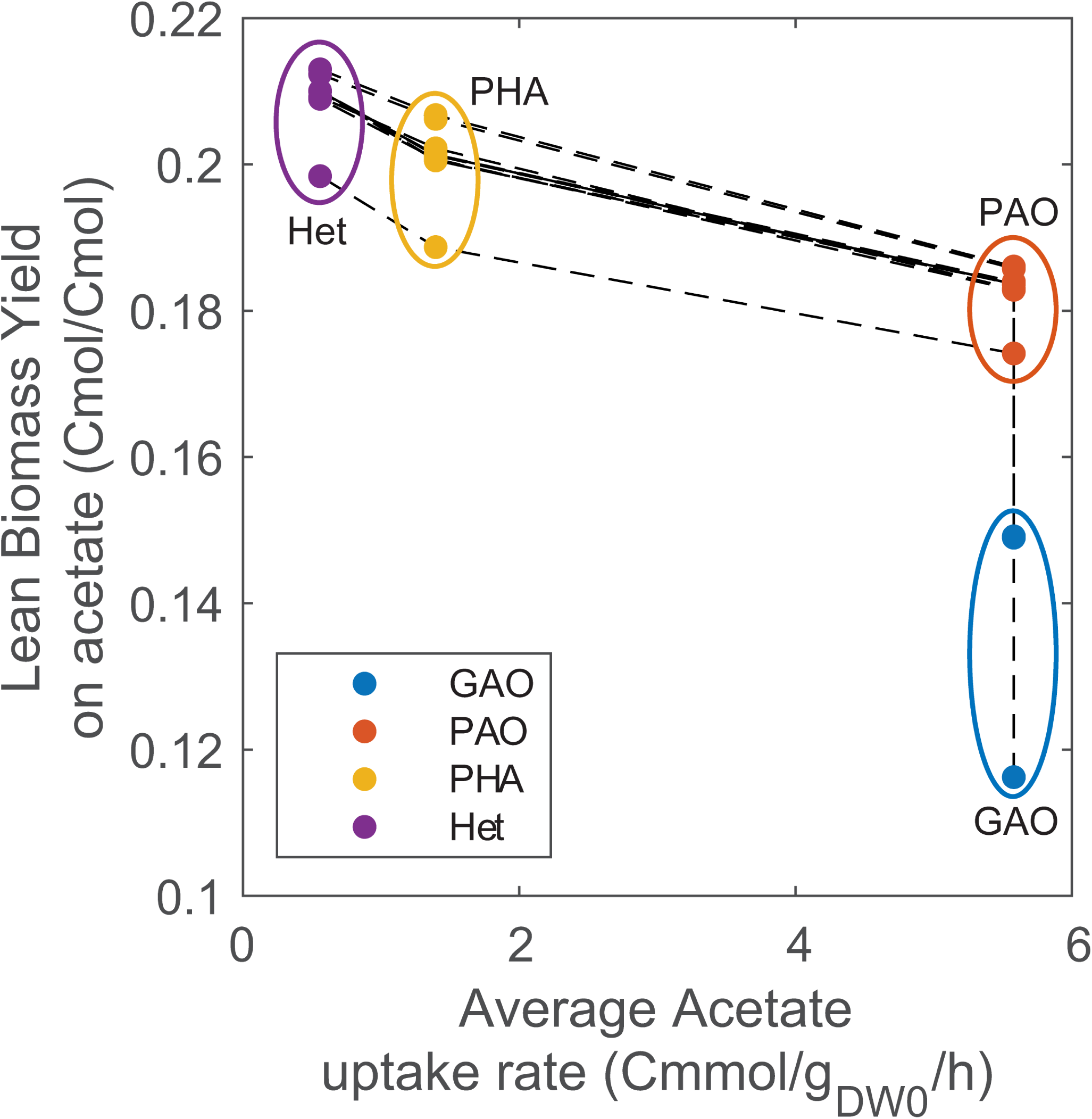
Rate/Yield trade-off curve for the sensitivity analysis of the turnover rates *(k*_cat_). The curve was obtained based on the simulated results of the different metabolic strategies shown in Figure 8 and the sensitivity analysis for GAO. The output from the simulation of the decrease of the *k*_*cat*_ of BMP synthesis was excluded from the figure.

From Figure 6 it becomes clear that the rate advantage that accumulating organisms (-AOs) have does not only come from fast acetate sequestration, but also from having the enzymes and resources that provide ATP anaerobically. To compare the different phenotypes (Figure 5) in a pareto-like relation between rate and yield (Figure 7), average acetate uptake rates were calculated taking into account the whole time in which acetate is available until it is fully consumed. For example, PHA-AOs only take 30 min to consume acetate but they only do so after the 1.5 h of anaerobic phase, thus the average rate is calculated over the 2 h of acetate availability.

In systems where carbon sources are provided all-at-once and are in excess (i.e. in batch cultivation), a rate-strategy is likely winning. With the current information provided to the model, GAOs and PAOs are equally capable of quickly taking up acetate anaerobically and to store it in the form of PHA. However, it is to be expected that in a mixed population of PAOs and GAOs, if each takes an equal share of the substrate, PAOs will be more efficient and thus create more offspring. The competition between PAO and GAO has been extensively studied and there are known environmental factors which balance the competition to either the PAO or GAO side; Changes in pH affect the ATP stoichiometry of transport processes, changes in temperature have a direct impact on the kinetic capacities of pathways and, as demonstrated earlier, reduced phosphate availability can have a detrimental effect on polyphosphate-based metabolism (Acevedo et al., 2012; Lopez-Vazquez et al., 2009; Smolders et al., 1994a; Welles et al., 2015). Here, we propose taking into account proton (H^+^) balancing between the intra- and extracellular space and proton translocation energetic costs in future simulations as a differentiator between PAOs and GAOs.

Another possible metabolic strategy to take up acetate anaerobically is employed by methanogens. While the acetate uptake rate is comparable, (about 5-7 Cmmol/gDW/h as reported for *Methanothrix spp.* by (J.W.H., Elferink, Visser, Hulshoff Pol, & Stams, 1994)), the biomass yield is not comparable as methanogens excrete the majority of the energy present in acetate in the form of methane, thus only harvesting a small fraction for their biomass.

In the scenario where no acetate is taken up anaerobically or when it is pulse-fed into an aerobic system, PHA-AOs are known to thrive and dominate in such conditions (Beun, Paletta, Van Loosdrecht, & Heijnen, 2000; Van Loosdrecht et al., 1997). Here, and again, organisms capable of sequestering substrate quickly will have a competitive advantage over yield-strategists. Finally, when PAOs, GAOs, and PHA-AOs fail at quickly sequestering substrate because of a) limitation of another nutrient, or b) inhibitors/predators, or c) when substrates are not readily available and need to be transformed first, or d) when substrates are continuously fed and limiting (i.e. in a chemostat), only then aerobic heterotrophs thrive. It is also important to note that simulations predicted that the most efficient strategy for PHA-AOs and heterotrophs would be to make use of glycogen and polyphosphate storage to cover for maintenance costs during the anaerobic starvation period. However, they may trade-off this (simulated) robustness to have a higher biomass yield. This will result in higher cell death during the anaerobic starvation period compensated by having higher amounts of offspring aerobically. In order to evaluate the investments that such strategy requires, enzyme and biomass decay should also be considered in future simulations.

### A tool to predict phenotypes based on defined environment

The new modeling framework developed by (Rügen et al., 2015) was used in this study as it allows for understanding growth in a periodic environment. Here we show how it can be applied to microbial communities by using a meta-network of the main functional groups present in such community and how it can predict which metabolic trait (phenotype) has the highest fitness for the set environment (i.e. combination of pathways that lead to the optimal energy allocation strategy).

Apart from using a meta-network instead of pre-defined strain-specific stoichiometric models, this formalism only requires a rough estimation of kinetic parameters to simulate an approximate proteome allocation. This approach contrasts with current kinetic models for PAOs/GAOs/PHA-AOs that use pre-defined optimal metabolic strategies, for example (Acevedo, Borrás, Oehmen, & Barat, 2014; Lanham et al., 2014; Marang, van Loosdrecht, & Kleerebezem, 2015). These kinetic models require extensive re-calibration to simulate different environments and consequently cannot predict the outcome of a new environmental condition. Furthermore, they may be used for describing microbial competition among pre-defined metabolic strategies (input), however they will not be able to predict the most optimal and competitive strategy (output). For example, it is not trivial that to maximize biomass in a competitive system, investment in PHA cycling is what enables fast substrate sequestration that in turn results in more offspring than a biomass yield-strategy.

This set of simulations has demonstrated why a polyphosphate-based metabolism is most adequate to cope with a competitive environment where oxygen is periodically unavailable. However, when polyphosphate is limiting, a glycogen-based metabolism emerges as an alternative. Lastly, simulations show that a faster and anaerobic acetate uptake is a strategic investment on competitiveness and represents a clear trade-off against growth.

The balancing between rate versus yield strategies has regularly been shown in batch and chemostat studies (Bachmann, Molenaar, Branco dos Santos, & Teusink, 2017). Here we showed that it is also the driver in microbial selection for dynamic ecosystems like wastewater treatment processes, but likely also tidal zones and systems subjected to light/dark cycles.

This is an example of when *“Unity in biochemistry”* (Kluyver & Donker, 1926) meets *“Everything is everywhere, but the environment selects”* (Baas-Becking, 1934): one metabolic network was used for all metabolic strategies (phenotypes) mentioned but it is the environment that sets which phenotype will thrive in the end.

## MATERIALS & METHODS

### Simulation and implementation

The files for the simulation were obtained from (Rügen et al., 2015) and executed in MATLAB version 9.4 (R2018a). The following changes were implemented: 1) LINPROG was used as linear optimization solver; 2) the simulation was adapted to this case study to model the usage of a defined external carbon source (acetate), within a specified amount of time. To this effect, acetate is modelled as an unbalanced metabolite (Ac_ext), however without any contribution to the composition of biomass (i.e. it has a weight of 0 on the Biomass composition vector). A spontaneous reaction (Ac_feed) was added to ensure the replenishment of acetate for the next cycle, which is only active at the end of the cycle.

All simulations were performed to mimic the experimental set-up used by (Acevedo et al., 2012), i.e. cycles of 5 hours divided into 20 uniform intervals of 15 minutes, with 1.5 hours of anaerobiosis and 3.5 hours of aerobic phase. To represent these two different conditions, ETC was set to 0 (i.e. blocked) and redTCA as irreversible during anaerobiosis, and then both were let free during the aerobic phase.

To simulate the different scenarios, the following constraints were used:

- Polyphosphate production and consumption reactions were set to zero (i.e. blocked) to obtain the results presented in Table 2.
- The maximal time for acetate uptake was constrained to 30 min, 1h30, 2h and 5h to produce Figure 5. For the remaining simulations, this value is set to 30 min.

## DATA AVAILABILITY

- Original model available as supplementary information of (Rügen et al., 2015)
- Adapted model available at https://github.com/cell-systems-engineering-tud/energy-allocation.git
- Experimental data available at (Acevedo et al., 2012)

## ACKNOWLEDGEMENTS

These investigations were supported by the SIAM Gravitation Grant 024.002.002, the Netherlands Organization for Scientific Research (NWO). The authors would like to thank Professor J. J. (Sef) Heijnen for his invaluable advice on this manuscript.

## AUTHOR CONTRIBUTIONS

LGdS, STM and SAW conceived the project. LGdS and STM performed computational analyses and wrote the manuscript with the input from the other authors. All authors read and approved the final manuscript.

## SUPPLEMENTARY INFORMATION

### Sensitivity analysis - Turnover rate, *k*_*cat*_

In order to study the effect of the value of the different turnover rates, *k*_*cat*_, a sensitivity analysis was made by varying each assumed *k*_*cat*_ one order of magnitude higher/lower. As it can be seen in Figure 8, the change in these parameters does not alter the qualitative analysis performed in this study, except for the decrease of *k*_*cat*_ for BMP synthesis. However, that simulation leads to about one order of magnitude lower biomass yield as compared with experimental data (Table 2), which indicates the unlikelihood of this value.

All these simulations, together with the sensitivity analysis for the GAM were used to generate Figure 9, showing that the conclusions drawn from Figure 7 remain the same.

### Full or partial operation of the TCA cycle

The usage of a full TCA cycle anaerobically has been hypothesized many times in PAO literature (Lemos, Serafim, Santos, Reis, & Santos, 2003; Louie, Mah, Oldham, & Ramey, 2000; Pereira et al., 1996; Zhou, Pijuan, Zeng, & Yuan, 2009). In this model, the full TCA cycle can be set active in the anaerobic phase by unbounding redTCA during both anaerobic and aerobic phases. In this scenario (Figure 10b), less glycogen cycling is needed compared to the reference simulation (Figure 4 and Figure 10a). These simulated results agree with the simulations done earlier in (Silva et al., 2018), in which it is discussed that a full operation of the TCA cycle anaerobically could explain the higher PHA yields found in EBPR than predicted by models. A higher PHA yield corresponds to a higher biomass yield, thus a fully operating TCA cycle in the absence of an external electron acceptor represents an improved polyphosphate-based metabolism (PAM).

**Figure 10.**
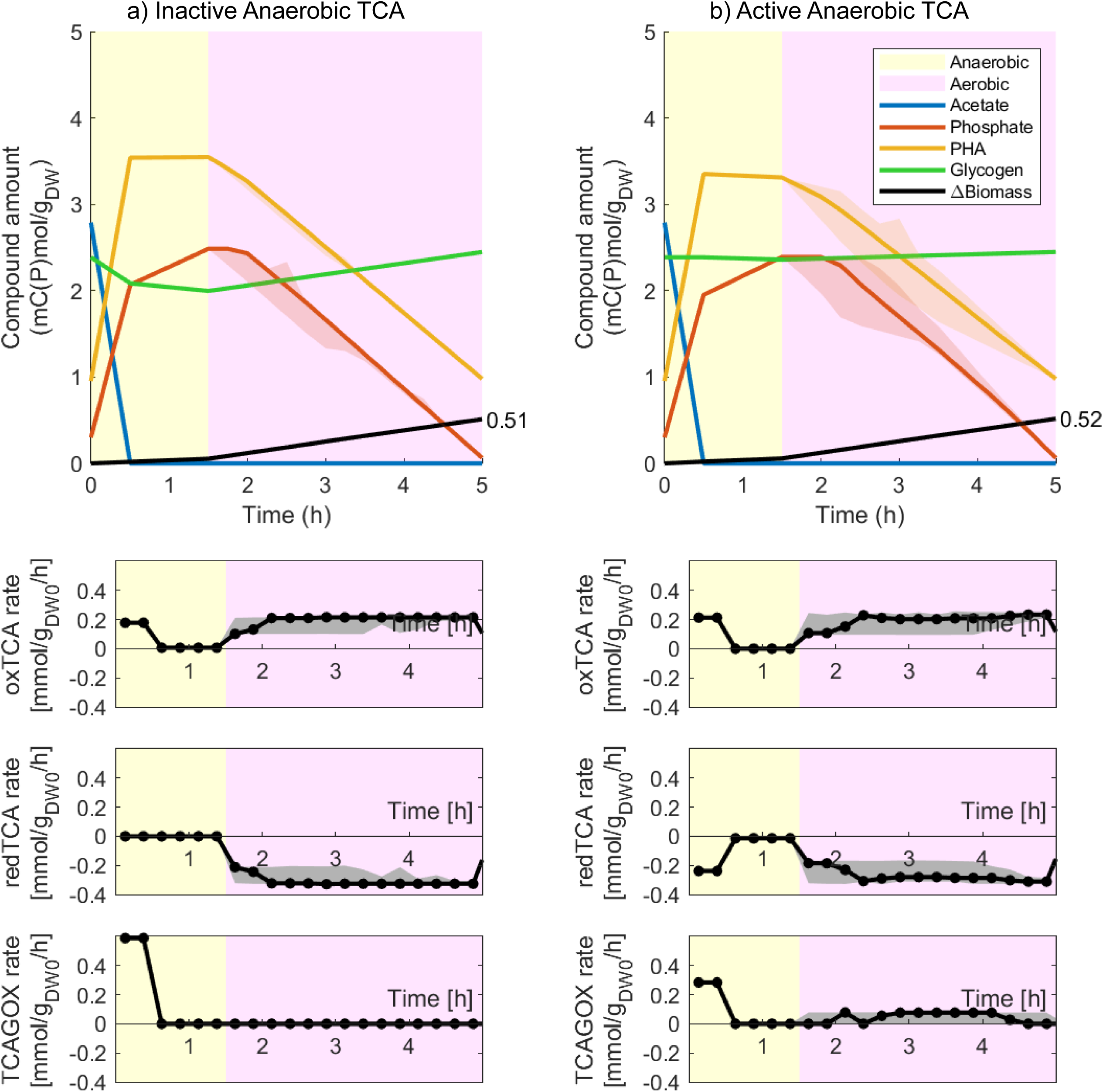
Simulations comparing the impact of an (in)active TCA cycle in the absence of an external electron acceptor (anaerobic): a) Reference simulation with an inactive TCA cycle anaerobically and b) with an active anaerobic TCA cycle.

When the TCA cycle is blocked at the level of the succinate dehydrogenase (Figure 10a), then a combination of glycogen metabolism with the oxidative TCA branch (isocitrate to succinyl-CoA) and the glyoxylate shunt are used. This latter strategy yields 73% PHB/PHA in a Cmol basis, while when the TCA is fully active, 90% of all PHA is PHB.

This change in strategy also impacts the biomass yield in each case being the best strategy when the TCA cycle is fully operational anaerobically (0.19 instead of 0.18 Cmol biomass per Cmol acetate). One of the simplifications in this model is to consider all reducing equivalents are in the form of NADH however, FADH is produced in the succinate dehydrogenase reaction step. The mechanisms for re-oxidizing this FADH, even if feasible without an external electron acceptor, may come with additional ATP costs, which should be considered in a future simulation.

